# A molecular lid mechanism of K^+^ channel blocker action revealed by a cone peptide

**DOI:** 10.1101/2020.06.15.153577

**Authors:** Chandamita Saikia, Orly Dym, Hagit Altman-Gueta, Dalia Gordon, Eitan Reuveny, Izhar Karbat

## Abstract

Many venomous organisms carry in their arsenal short polypeptides that block K^+^ channels in a highly selective manner. These toxins may compete with the permeating ions directly via a “plug” mechanism or indirectly via a “pore-collapse” mechanism. An alternative “lid” mechanism was proposed but remained poorly defined. Here we study the block of the Drosophila Shaker channel by Conknunitzin-S1 and Conkunitzin-C3, two highly similar toxins derived from cone venom. Despite their similarity, the two peptides exhibited differences in their binding poses and in biophysical assays, implying discrete modes of action. We show that while Conknunitzin-S1 binds tightly to the channel turret and acts via a “pore-collapse” mechanism, Conkunitzin-C3 does not contact this region. Instead, Conk-C3 uses a non-conserved Arg to divert the permeant ions and trap them in off-axis cryptic sites above the SF, a mechanism we term a “molecular-lid”. Our study provides an atomic description of the “lid” K^+^ blocking mode and offers valuable insights for the design of therapeutics based on venom peptides.

## Introduction

Ion channels are pore-forming proteins integrated into the membranes of all cells. They allow the passage of specific populations of ions through the membrane, based on the electrochemical ion gradients across its two sides. Among the ion channels, potassium channels (K+) are the most abundant and diverse group, having crucial roles in a myriad of cellular processes, ranging from the generation of electrical impulses in nerve and muscle cells to hormone secretion and blood pressure regulation^1^. Most functional K^+^ channels are tetramers, where the four subunits assemble to form a water-filled pore that enables a rapid and selective translocation of K^+^ ions between the intracellular and the extracellular faces of the hydrophobic lipid bilayer^2^. Several constrictions along this aqueous path contribute the various filters and gates that regulate ion movement. The “Selectivity filter” (SF) is a structural element that is common to all known K^+^ channels and is highly conserved from bacteria to mammals^3^. It consists of a 5-residue signature sequence (TI/VGY/FG) contributed by each subunit that forms a tight constriction at the extracellular face of the pore. In most voltage-gated and inwardly rectifying K^+^ channels, a conserved Asp/Glu residue appears at the extracellular entry to the pore. To pass through the narrow selectivity filter, a K^+^ ion must first shed its hydration shell. This process is catalyzed by the SF signature residues, which introduce carbonyl oxygen atoms into the lumen of the pore, serving as a surrogate hydration shell and reducing the energetic penalty for the ion dehydration process^4,5^.

Many peptide blockers, isolated from the venoms of scorpions, bees, snakes, cone snails, and sea anemones antagonize the process of selective K^+^ permeation by a physical occlusion of the pore^6^. In most studied cases, a “cork in a bottle” or “plug” mechanism was identified, in which a conserved lysine sidechain of the toxin is inserted into the channel pore and interact with the uppermost ring of SF carbonyls, precluding K^+^ coordination^7–10^. Given the involvement of K^+^ channels in diverse pathological conditions such as cancer, autoimmune disease, and neurological/cardiac disorders, numerous efforts aiming to identify novel K^+^ peptide blockers or to improve the pharmacological profiles of existing ones, were made^11^. Building upon the foundations of the “molecular plug” mechanism, many of these studies aimed to enhance the potency and the specificity of a given toxin molecule by introducing various natural and chemical modifications, while maintaining the crucial interaction between the conserved lysine and the channel pore^12–17^.

Yet, many venom-derived peptide K^+^ blockers lack a conserved pore-inserting lysine or exhibit binding modes inconsistent with a “molecular plug” mechanism. A general mode of action referred to as “lid-block” or “turret-block” was proposed, whereby the toxin interferes with ion translocation by forming a lid above the channel pore^16,18,19^, however, a detailed molecular description for such mechanism is still lacking. Recently, we have described the mode of action of Conkunitzin-S1, a cone snail toxin that potently blocks the drosophila Shaker channel. We have found that instead of plugging the pore, the toxin triggers the collapse of the SF by targeting channel residues that gate the access of water molecules to the aqueous cavities around the pore, resembling the molecular events that take place during slow inactivation^20^.

Herein we describe the blocking mechanism of Conk-C3, a yet undescribed toxin isolated from the venom of *Conus Consors*, that exhibits 86% similarity to Conkunitzin-S1. To our surprise, despite this similarity, MD simulations of Conk-C3 predicted an alternative binding mode, which has not triggered a collapse of the pore. Instead, we observed that the toxin utilizes a non-conserved Arg sidechain to divert the permeant ions electrostatically and to trap them in off-axis cryptic sites above the SF. Predictions derived from these simulations were verified experimentally, and their molecular underpinnings were pinpointed using a chimeric approach.

The findings offer a molecular insight into the selective interactions of peptide blockers with K^+^ channels and suggest that computer-aided modeling may have reached the maturity level required for its use as a practical tool in the design of novel therapeutics using venom peptides as leads.

## Results

### Conk-C1 and Conc-C3 – novel toxins from Conus Consors with high potency at the Drosophila Shaker channel

Two transcripts encoding putative kunitz-like peptides were acquired during a proteomic analysis of the fish-hunting cone snail *Conus Consors* venom^21^ and designated Conkunitzin-C1 (Conk-C1) and Conkunitzin-C3 (Conk-C3). Both peptides exhibited high similarity to Conkunitzin-S1 (Conk-S1) from *Conus Striatus*, which was described previously^20,22^ (85% and 86% similarity for Conk-C1 and Conk-C3, respectively, Fig. 1A). Synthetic genes encoding for Conk-C1 and Conk-C3 were constructed, the peptides were expressed in E. coli and purified using an established method^20^. Both peptides were crystallized, and high-resolution structures were solved from x-ray diffraction, reveling a typical Kunitz fold composed of a double-stranded antiparallel β-sheet and N- and C-terminal α-helices reticulated by two disulfide bridges (Fig. 1B). Initial pharmacological profiling revealed minor activities of these peptides at mammalian K_v_ channels (K_v_1.1 to 1.3, <10% block at 5μM) but higher potencies at the drosophila *shaker* isoform (Fig. S1). Similar to Conk-S1^20^, Conk-C1 was significantly more potent at the Shaker K427D mutant, indicative of an interaction between the toxin and the channel turret region (Fig. S1). In contrast, the block of Shaker by Conk-C3 was insensitive to the K427D mutation, and the toxin had an affinity that is two orders of magnitude lower than that of Conk-S1 (190nM for Conk-C3 vs 1nM for Conk-S1, Fig. 2A). Based on these observations, we surmised that despite its high similarity to Conk-S1, Conk-3 might utilize a distinct binding mode to block the Shaker channel.

**Figure 1.**
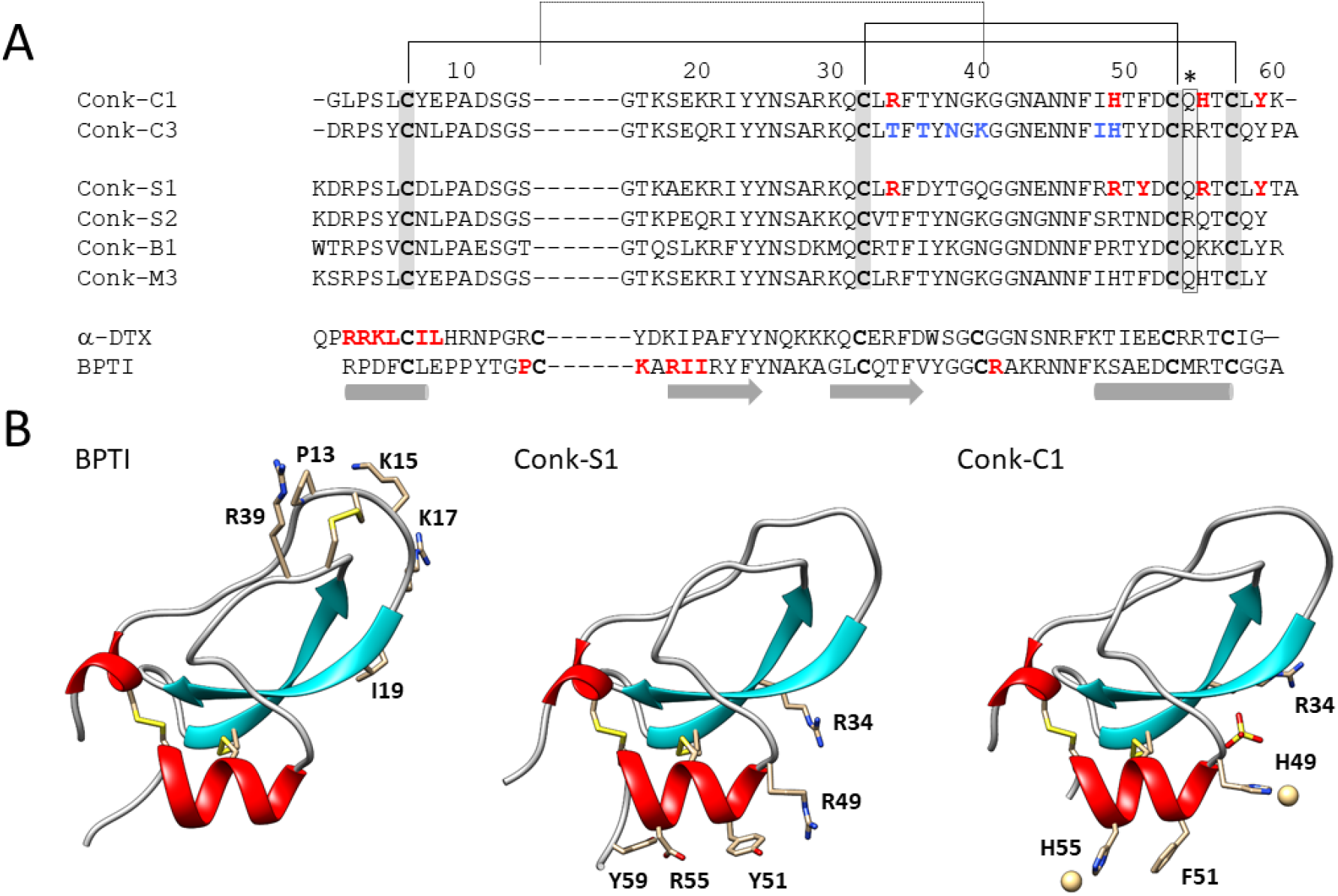
A, Sequence alignment of selected Kunitz-fold peptides. Conserved cysteine residues are shaded in grey. Two conserved disulfide bridges are indicated in solid lines. A third, non-conserved bridge is indicated by a dotted line. Secondary structure elements are depicted at the bottom. A non-conserved arginine identified herein as a key residue for channel block by Conk-C3 is indicated by asterisks. Conk-C1 and Conk-C3 are from *Conus Consors*, as described herein; Conk-S1 and Conk-S2 from Conus Striatus^22^; Conk-B1 is from Conus Bullatus^53^; Conk-M3 is from Pionoconus magus^54^; α-DTX is from *Dendroaspis angusticeps* snake venom^55^, and BPTI is from Bos Taurus^56^. The bioactive residues of Conk-S1, α-DTX, and BPTI are highlighted in red; for Conk-C3, positions that markedly differ from Conk-S1 are highlighted in blue. B, Ribbons representations of BPTI, Conk-S1, and Conk-C1. Disulfide bridges are in yellow sticks. For BPTI and Conk-S1, the bioactive residues, as highlighted in A, are shown. For Conk-C1, ions that have co-crystallized with the protein are depicted.

**Figure 2.**
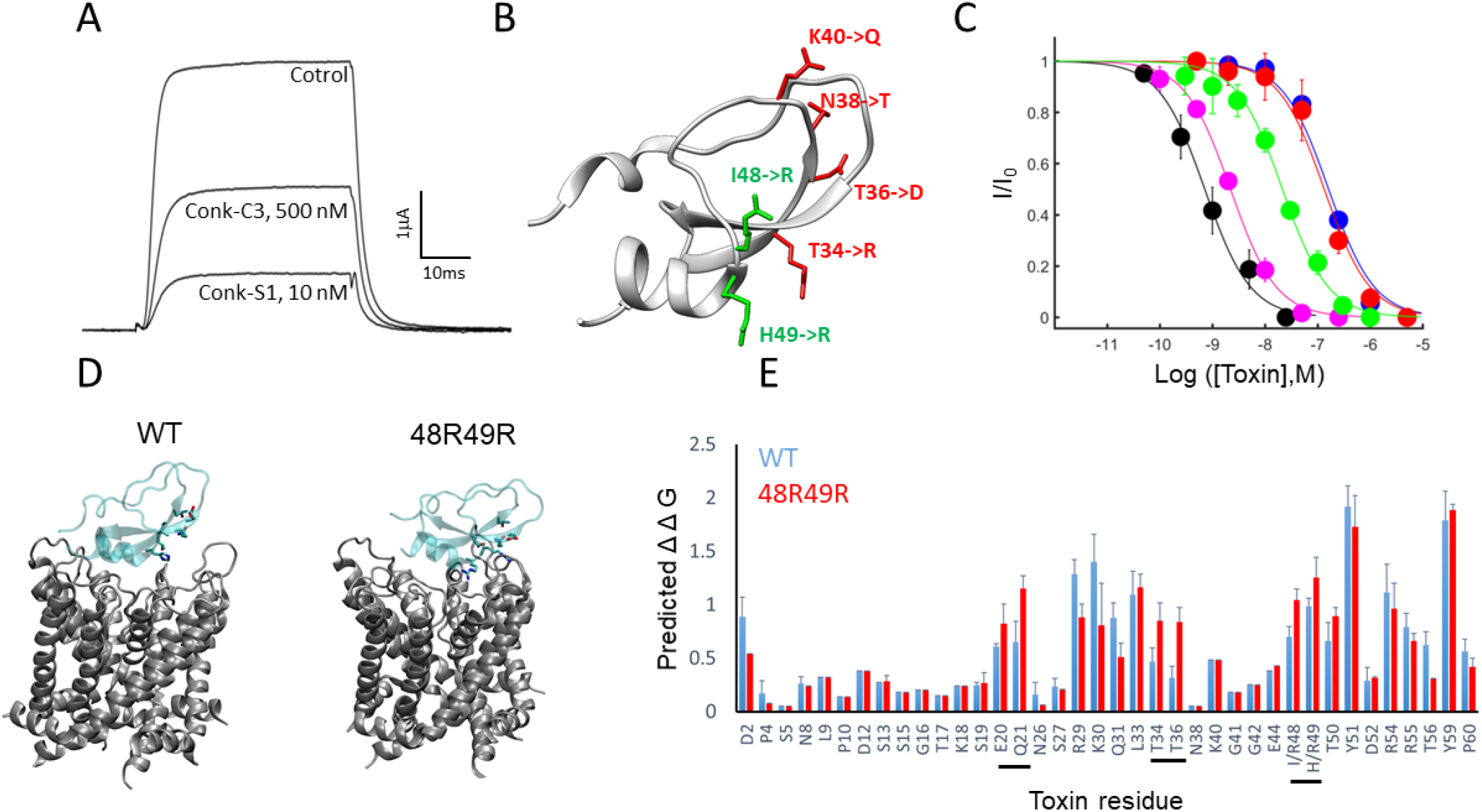
Reconstruction of Conk-S1 binding determinants on the scaffold of Conk-C3. A, Outward current traces recorded from an oocyte expressing Shaker_KD_ in response to 0mV step depolarization in the presence of 500nM Conk-C3, or 10nM Conk-S1. B, Substitutions performed on Conk-C3 backbone to produce the Conk-C3^RDTQ^ (red) and the Conk-C3^RR^ (green) chimeric toxins. C, Dose-response curves of Conk-S1 (black, 0.95±0.03nM), Conk-C3 (Blue, 162±10nM), Conk-C3^RDTQ^ (red, 132±22nM) and Conk-C3^RR^ (green, 2.7±1nM) on Shaker_KD_. D. Snapshots of Conk-C3 (left) and Conk-C3^RR^ (right) bound to Shaker_KD_ taken at the end of a 100ns production runs. The substituted residues at positions 48, 49, as well as residues that contact the turret of chain D in the model of Conk-C3^RR^, are highlighted. E, Computational alanine scan performed on the models depicted in D. The residues highlighted in D are underlined.

### High potency requires the coordinated binding of two discrete residue clusters

Analysis of Conk-S1 and Conk-C3 sequences revealed two clusters of residues that vary between the toxins. Residues R34, D36, T38, and Q40 residing at the second β-strand of Conk-S1 are substituted by T34, T36, N38, and K40 in Conk-C3; the arginine side chains at positions 48,49 of Conk-S1 are substituted by Ile and His in Conk-C3 (Fig. 2B). We have constructed chimeric peptides bearing the above Conk-S1 segments on the backbone of Conk-C3, aiming to assess the contribution of these differences to the variable potencies of the parental toxins at Shaker_KD_. While reconstitution of Conk-S1 second β-strand on the scaffold of Conk-C3 (Conk-C3^RDTQ^, 34R/36D/38T/40Q) had no effect on potency, the I48R/H49R substitution has increased potency 10-fold (Conk3^RR^, Fig. 2C). Curiously, we observed an additional increase in potency when these two substitutions were combined, resulting in a chimeric toxin with potency at the nanomolar range, reminiscent of Conk-S1 (Fig. 2C). Thus, it seems that both segments contribute to the high potency of Conk-S1 at Shaker_KD_ and that the contribution of the β-strand residues requires a prior adjustment near the C-terminal α-helical segment of the toxin.

### MD simulations reveal distinct binding modes for Conk-3 and Conk-3^RR^

To gain structural insight into these experimental observations, we docked Conk-C3 and the Conk-C3^RR^ mutant onto Shaker_KD_, and employed unbiased molecular dynamics to relax any constraints and refine the initial docked models. Both toxin species assumed distinct binding poses within the initial 20ns segment of the MD run, which remained stable for the duration of the trajectory, up to 0.5μs. Snapshots taken at t=100ns from simulations of bound Conk-C3^RR^ were highly homogenous (rmsd = 1.34±0.2Å over all backbone atoms, 9 simulations), while models of wt Conk-C3 exhibited heterogeneity (rmsd = 2.53±0.54Å, 14 simulations). Clustering of the Conk-C3 wt models revealed a large homogenous sub-population accounting for 60% of the runs (rmsd = 2.12±0.13Å, 8 out of 14 runs), that were selected for further analysis.

Curiously, the stable binding pose of the Conk-C3^RR^ mutant was tilted by 12° compared to that of the wild type toxin, resulting in an improved packing of the toxin molecule against the turret region of subunit D of the channel (Fig. 2D). Computational alanine scanning predicted higher contributions of the β-sheet residues Q21, T34 and T36 to the free energy of binding, resulting from multiple polar interactions formed between these residues and the channel turret (Fig. 2E, S2A). The newly introduced R49 is predicted to form a stable cation-π interaction with F425^D^, and a salt bridge / polar interactions with the nearby D447^D^ and T449^C^, respectively (Fig. S2B). In stark contrast, in MD trajectories of the wild-type Conk-C3 complex, H49 that occupies the equivalent position forms an intramolecular salt-bridge with the nearby D52 and does not initially contribute any stable contacts with the channel protein (Fig. S2C). Instead, D447^D^ is often found in association with R54, a residue that is non-conserved in other conkunitzins (Fig. 1A). In addition, In Conk3^RR^, The sidechain of Y59 penetrates deeper into the peripheral cavity of chain B, where it makes an aromatic interaction with F425^B^, and coordinates via hydrogen bonds the sidechains of D447^B^ and T449^A^ (Fig. S2D). These differences in contacts made by the two toxin species with the channel protein profoundly altered their simulated effects on the solute molecules and on the permeant ions, as described below.

### *Conk-3^RR^*, but not Conk-C3, induce an asymmetric collapse of the SF

MD simulations performed with unmodified K^+^ channels revealed remarkable stability of the selectivity filter region, which typically maintains its symmetrical conformation and bound ions up to several microseconds^20,23,24^. Recently we have demonstrated that Conk-S1, which highly resembles Conk-C3, could trigger an asymmetric collapse of the channel pore by introducing ectopic traffic at the water-filled cavities that surround the channel pore^20^. Infiltration of water from these cavities into the SF can then displace the bound ions and promote pore collapse. This effect was mediated by interactions formed between the toxin and the ring of D447 residues above the SF, which gate the water flow in the peripheral cavities via hydrogen bonds with W434(‘DW-Gates’^20^). With this mechanism in mind, we turned to probe the filter-symmetry in MD simulations of the Shaker-Conk-C3 complex. As opposed to Conk-S1, in simulations of Conk-C3 bound ions at the SF as well as the symmetry of the pore were invariably retained (Fig. 3A, B). Slightly asymmetric conformations (Δ(B-D,A-C)>1Å) were observed in 3 out of the 8 runs, resulting from slight dilation at the upper section of the SF (Fig. S3A). In contrast, in 6 out of 9 simulations carried in the presence of the 48R49R mutant, we observed loss of the ion bound at S2 during the initial 20ns equilibration phase or early in the subsequent production run, followed by an asymmetric constriction of the SF (Δ(B-D,A-C) > 2Å, Fig. 3A, S3B-E). We have found clues to the molecular underpinnings of this striking difference in the behavior of the two systems by examining the water traffic at the peripheral cavities (Fig. 3C). In both systems, we observed a significant increase in the flow of water molecules at the peripheral cavity of chain D as compared to control runs carried without a bound toxin (Fig. 3C). These observations were correlated with predominantly open states of the DW gates in chain D (DW^D^) in the presence of both toxins (Fig. 3D). For Conk-C3, the open state was stabilized by polar interactions formed between D447^D^ and R54/T50 of the toxin (Fig. S2D), while for Conk-C3^RR^ alternating interactions between D447^D^ and R49, T50, Y51, and R54 of the toxin were observed (Fig. S2B). In contrast, the effects of the bound toxins on the water traffic at the peripheral cavities of chain B differed considerably. While the wild-type toxin caused a ~4-fold reduction in water flow through DW^B^, Conk-C3^RR^ practically halted water exchange through this gate (Fig.3C). The deeper penetration of Conk-C3^RR^ Y59 towards the peripheral cavity Conk-C3^RR^ in chain B (Fig. S2B, D) invariably trapped DW^B^ in a locked conformation (< 3Å), as opposed to predominantly open conformations of this gate observed in simulations of the wild-type toxin (>4 Å, Fig. 3D). Due to these differences, the overall imbalance introduced to the water traffic around the channel pore by Conk-C3^RR^ was significantly higher compared to that induced wild-type toxin (Fig. 3C), which may rationalize why collapse of the channel pore was observed in presence of the former but not of the later^18^.

**Figure 3.**
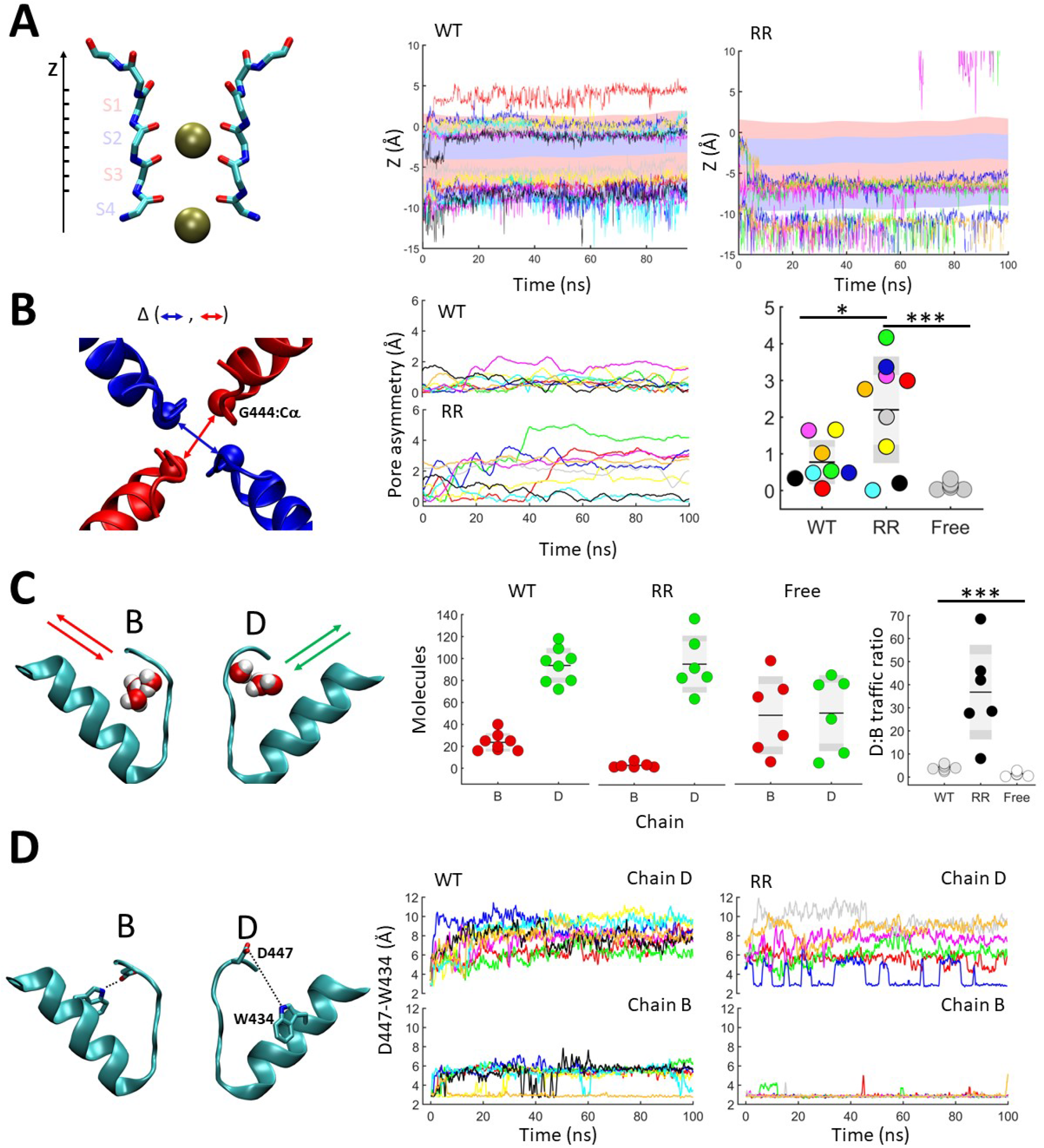
*Conk-3^RR^*, but not Conk-C3, induce an asymmetric collapse of the SF. Data were collected from eight simulations of Conk-C3, nine simulations of Conk-C3^RR^, and six control simulations carried with a free channel protein. The left pane in each row illustrates the quantified parameter. A, Vertical coordinates of K^+^ ions in the SF in Conk-C3 (left) and Conk-C3^RR^ (right) along the MD trajectory. Alternating red-blue stripes at the background delineate the s1 to s4 ion coordination sites. Each run is represented by a unique color that is maintained throughout the figure. B, Pore asymmetry measured as the difference between the cross-subunit diagonals (taken at the Cα atoms of G444) of chains B, D and chains A, C. middle: evolution of pore-asymmetry during trajectories of Conk-C3 (top) and Conk-C3^RR^ (bottom). Right: the pore asymmetries taken at t=100ns are depicted for the two toxin species and for free Shaker channels. C, Simulations of Conk-C3^RR^ exhibit highly asymmetric water traffic at the peripheral cavities of chains B and D. middle: each point represent the count of unique water molecules that were exchanged between the peripheral cavity and the bulk during a 100ns simulation window. For Conk-3^RR^, the displayed data were restricted to the six trajectories in which pore-collapse was observed. Right: the ratio between the number of unique water molecules exchanged between the peripheral pockets in chains D and B, and the bulk is presented for the three simulation systems. D, conformational dynamics of the DW gates in chains C and D of the Conk-C3 and Conk-C3^RR^ systems. The plots depict the distance between the Oδ1/Oδ2 atoms of Asp447 and the Nε1 atom of Trp434 in chains B and D along the MD trajectories.

### MD simulations suggest a molecular lid mechanism for Conk-C3

Unbiased MD simulations of bound Conk-C3 negated a pore-collapse mechanism for this toxin but offered little insight into alternative mechanisms. While we could detect transient interactions between R54 of the toxin and the backbone carbonyls of Y445 at the SF, in most trajectories R54 side chain assumed a more external position and formed salt bridges with D447 (Fig. S2D, 4A), negating its role as a “plug”. Moreover, we detected considerable water exchange between the outer vestibule of the channel pore and the bulk in the presence of the bound toxin, in stark difference from the classical pore-blockers CTX and Shk, which formed a tight seal around the channel pore (Fig. S4A).

If Conk-C3 does not “seal” nor “plug”, nor “collapse” the pore, how does it block the flow of ions?

To this end, we have employed steered MD to force the permeant ions against the bound toxin molecule. We carried seven simulations in which the toxin-bound system was allowed to equilibrate for 100ns before a 0.4V electric field directed outward was applied for an additional 100ns. During these simulations, we observed three types of interactions between the toxin and K^+^ ions bound at the SF, which often occurred consecutively along the trajectory as the ions progressively assumed more “outward” coordinates.

First, in cases in which R54 of the toxin has pointed into the pore at the end of the equilibration run, thus partially occupying the S0 K^+^ coordination site, the applied electric field induced a clear repulsion between this side chain and a permeant ion. K^+^ translocation from S2 to S0 triggered a rotation of R54, breaking its contacts with the backbone carbonyls of the SF while maintaining its interaction with D447 (Fig. 4A, Movie S1).

**Figure 4.**
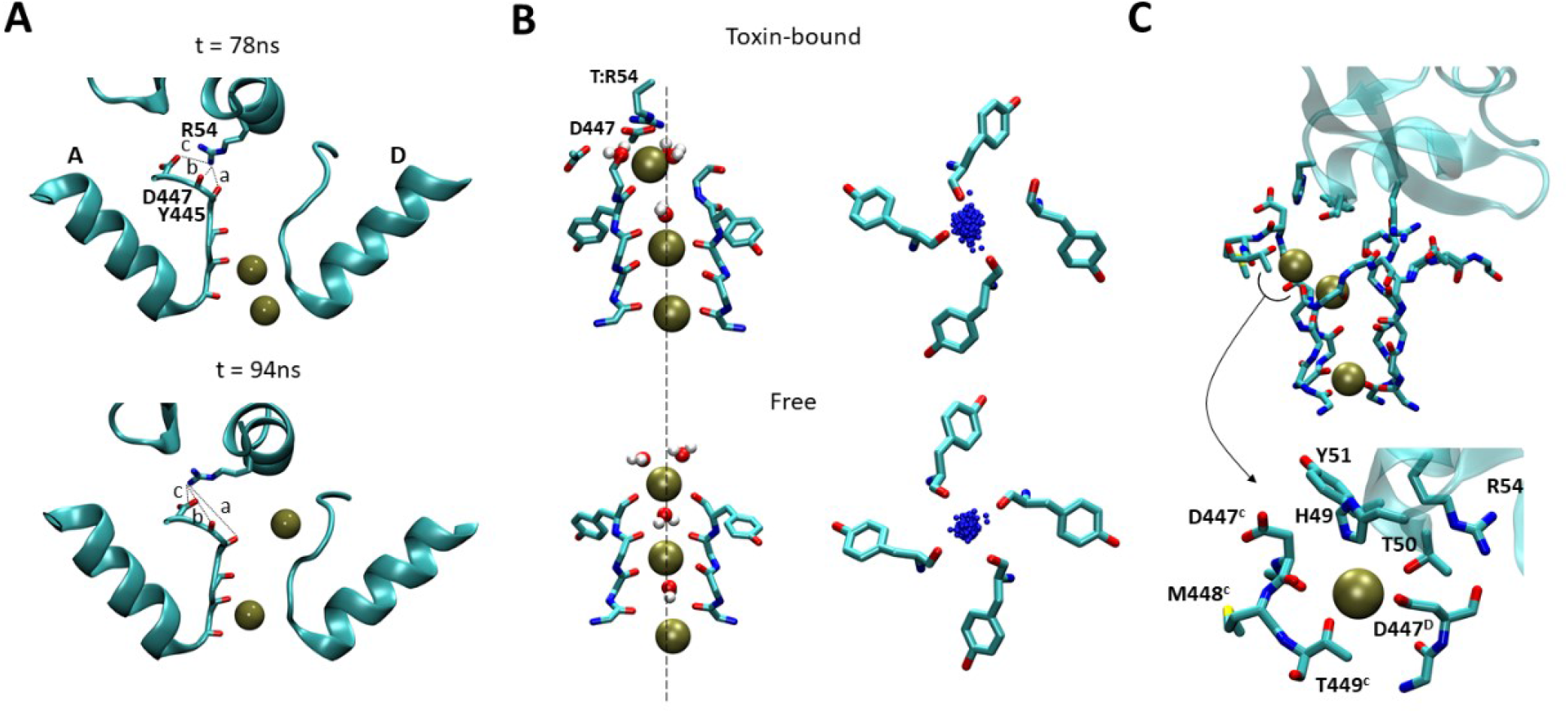
MD simulations suggest a molecular lid mechanism for Conk-C3. A, electrostatic repulsion between R54 and a K^+^ ion upon S2->S0 transition. The images depict snapshots of the channel selectivity filter and the bound toxin taken 78ns (top) and 94ns (bottom) following the application of a +0.4V electric field. Interactions made by the R54 side-chain are indicated. Measured distances were a=2.7Å, b=3.6Å and c=4Å at t=78ns and a =7.8Å, b = 4.6Å, c = 2.9Å at t=94ns. See also movie S1. B, asymmetric ion coordination at site S0. Drawing to the left depict the ionic configuration at the SF in snapshots taken during steered-MD simulations of Conk-C3 (top) and toxin-free (bottom) systems. The pore axis is indicated as a dashed line. Conk-C3:R54 and D447^A,D^ are rendered in sticks. Drawings to the right compare the spatial distributions of a K^+^ ion bound at the S0 site during a 50ns segments of toxin bound (top) and control (bottom) simulations. Tyr445 residues that contribute the bottom coordination ring of S0 are rendered in sticks. C, A cryptic K^+^ coordination site formed at the interface between chains C and D of the channel and the toxin molecule. Top: side view of the SF region following a second S2->S0 permeation transition. Bottom: zoom on the coordination site highlighting residues that contribute main-chain and side-chain oxygen atoms to coordinate the ion.

Second, Following S2->S0 transition, the ion bound at S0 was slightly diverted off the pore axis and away from T:R54 (Fig. 4B, S4B, C). This ionic configuration of the pore typically lasted for several tens to several hundreds of nanoseconds and was accompanied invariably by a gradual dilation of the upper segment of the pore, observed in all simulations carried with an applied electric field (Fig. S4B-E). We did not observe such dilation in control simulations conducted with Conk-C3^RR^ bound systems, in which ion access to S0 was blocked due to the SF constriction at the A-C plane (Fig. S4D-F).

Third, Following S2->S0 transition, under the persistent effect of the electric field, an incoming K^+^ ion could trigger an additional knock-off event, diverting the ion bound at S0 into a coordination pocket formed above the SF by backbone and sidechain atoms contributed by both the channel and toxin molecules (Fig. 4C). This hybrid ion-coordination site was partially shielded from the external solute by T50, H49, and Y51 of the toxin and persisted, depending on the initial configurations of these sidechains, from several nanoseconds to several tens of nanoseconds, after which the ion diffused to the bulk (movie S2).

Thus, MD simulations suggest that Conk-C3 obscure ion flow by a mixed mechanism in which a non-conserved arginine sidechain is utilized to divert the permeant ions off the pore axis and trap them in cryptic coordination sites formed at the toxin-channel interface.

### Experimental validation of predictions derived from the “molecular lid” model

The intricate atomic details offered by MD simulations can rarely be fully validated by experiment, owing to the sampling and the accuracy problems associated with MD on the one hand and to the limitations imposed by traditional biochemical/biophysical techniques on the other^25^. Yet, in some cases, mechanistic insights offered by MD simulations are very useful for providing non-trivial predictions that fuel experimental design. Subsequent iterations between simulation and experiment can be used to partially bridge the gap between the *in-silico* world to the “real” world^26^.

One such non-trivial prediction offered by MD simulations of K^+^ channel blockers relies on their different levels of interaction with the permeant ions. Classical pore blockers acting via a “molecular plug” mechanism directly compete with ions bound at the S1/S0 coordination sites, thus readily dissociate from their binding site when the external [K^+^] is raised, or a strong depolarization is applied^27,28^. In contrast, Conk-S1, predicted to block ion conduction by triggering the collapse of the pore, thus avoiding any interaction with the permeant ions, was resilient to voltage-induced dissociation^20^. The “molecular lid” mechanism described herein for Conk-C3 lies between these two extremes: while bound toxin did not forbid ion occupation at the upper coordination sites of the SF, antagonistic interactions, which seem to originate from electrostatic repulsion between the toxin and these ions, were observed (Fig. 4). In concert with this description, the behavior of Conk-C3 in depolarization-induced dissociation assays was intermediate between that of the classical blockers to that of Conk-S1 (Fig. 5). Strong depolarizations could only partially dissociate the bound toxin, and longer durations were required to saturate this effect as compared to the classical pore blocker CTX (Fig. 5B, C). The 48R49R mutation completely abolished toxin sensitivity to voltage-induced dissociation, consistent with the simulated changes in its binding mode and blocking mechanism (Fig. 5A).

**Figure 5.**
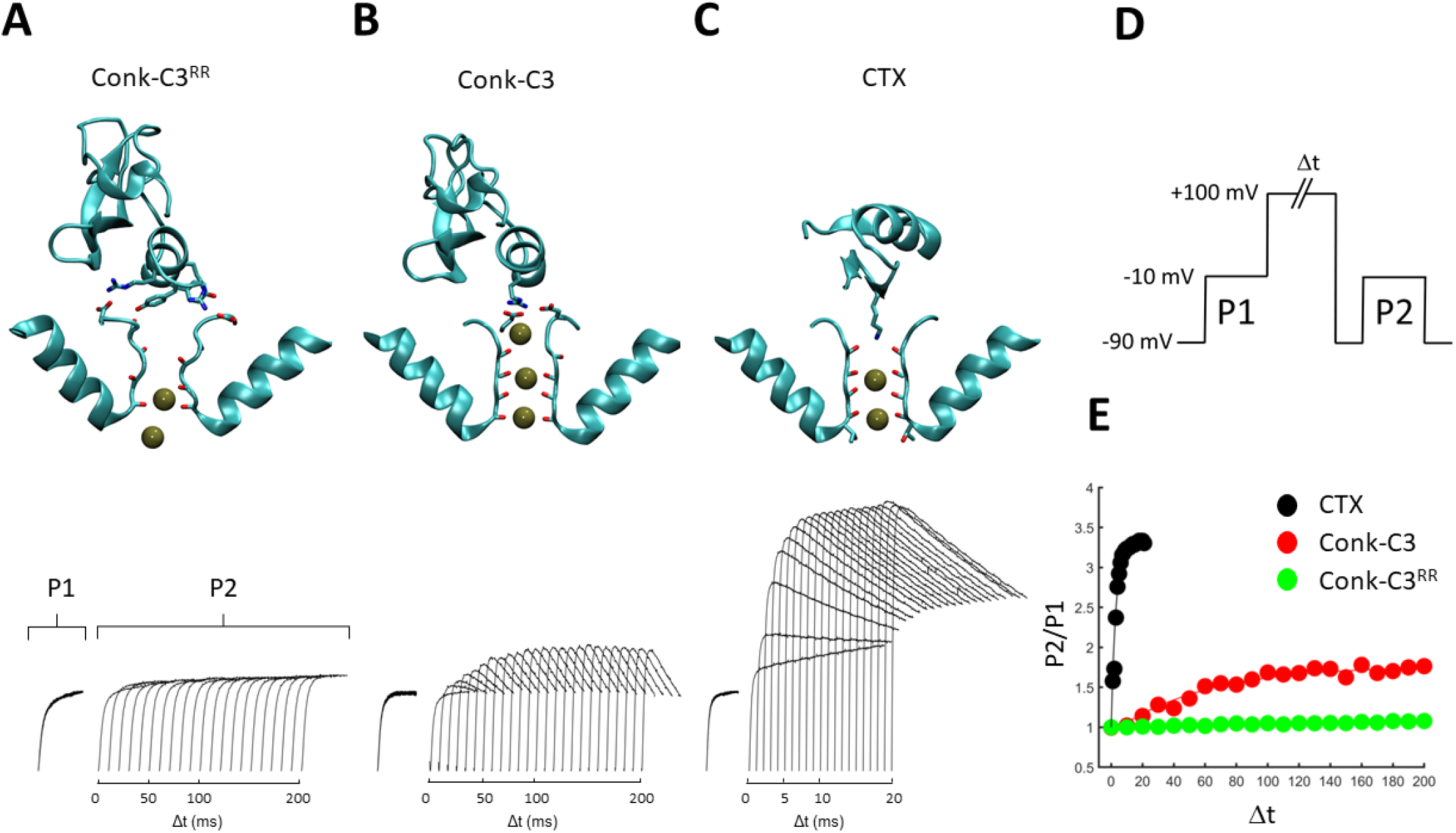
Trans-enhanced dissociation experiments are consistent with three distinct levels of interaction between peptide blockers and K^+^ ions in the selectivity filter. Top: snapshots taken at the ends of simulations carried with a 0.4V electric field applied. Two-opposing subunits with ions bound at the SF as well as toxin residues found critical for block by each toxin are displayed. Bottom: Typical currents recorded during P2 as a function of Δt in the presence of each toxin, during application of the voltage protocol depicted in D. P2 Currents from individual traces are overlaid using an arbitrary horizontal displacement, Δt is the length of the depolarizing pulse preceding P2. E, The ratio P2/P1 is plotted as a function of Δt for each toxin.

Additional prediction derived from the simulated difference in the binding modes of Conk-C3 and the Conk-C3^RR^ mutant is that the stronger interaction of the later with the turret of chain D should increase its sensitivity to alterations in this region. This prediction was met when we initially observed that Conk-C3^RR^, but not the wild-type toxin, is sensitive to the K427D substitution (Fig. 2). Further probing the contribution of turret residues to toxin binding revealed that a D431R substitution that had a deleterious effect on Conk-S1 binding had an only marginal impact on the binding of Conk-C3 (Fig. 6A). We additionally spotted a difference in the sensitivity of the two toxins to an M448K substitution in the channel, further supporting the proposed difference in their binding mode (Fig. 6A).

**Figure 6.**
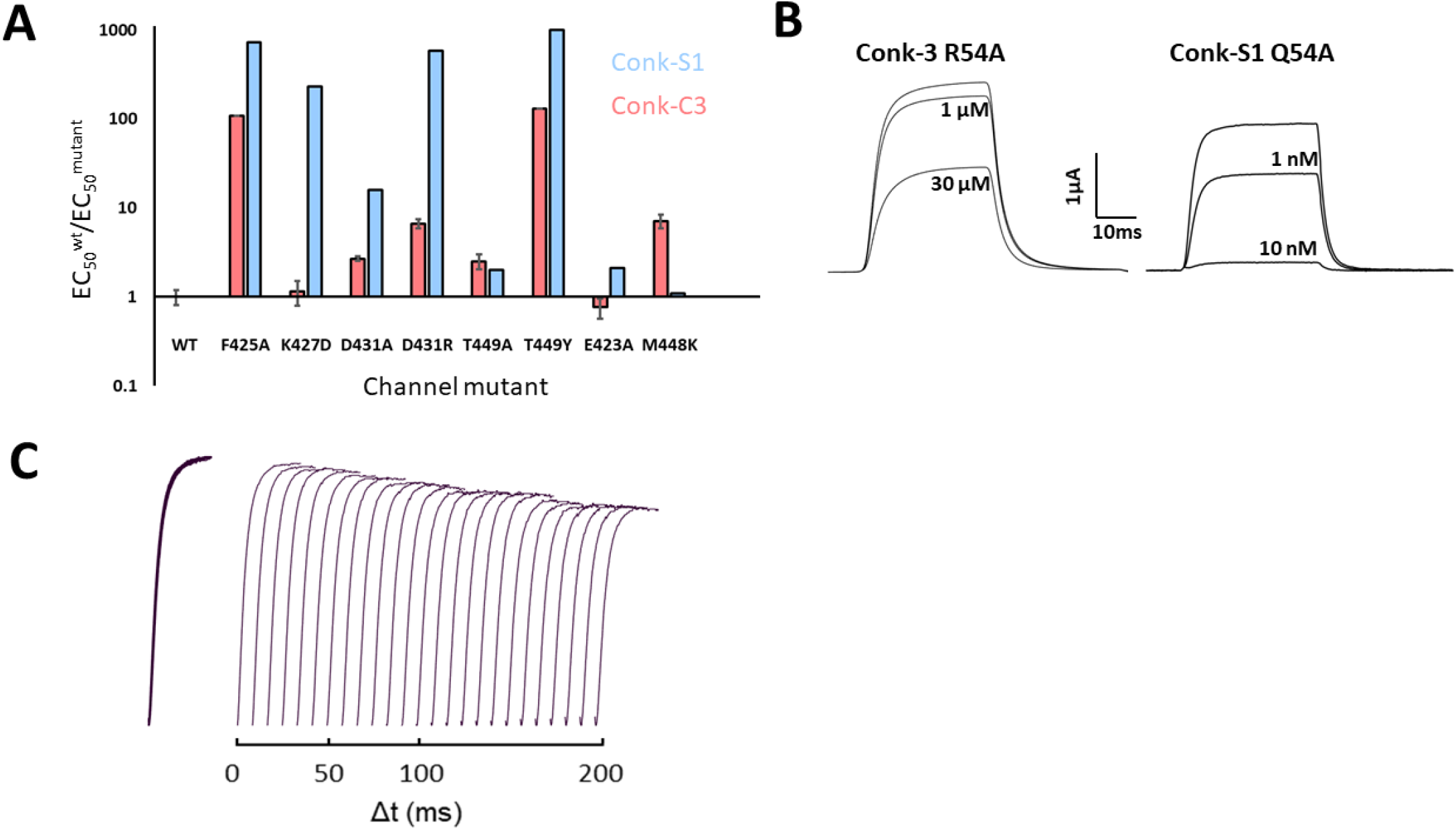
Experimental validation of predictions derived from the “molecular lid” model. A, the impacts of point mutations at Shaker_KD_ on Conk-S1 (blue) and Conk-C3 (red) are compared. Some of the Conk-S1 data were previously reported^20^. Each bar represent measurements from at least 3 oocytes. B, Traces recorded from Shaker_KD_ channels in the presence of Conk-C3 R54A (left) or Conk-S1 Q54A (right). C, Traces recorded during a trance-enhanced dissociation assay carried in the presence of 30μM Conk-C3. Protocol details were as in Fig. 5.

Finally, and perhaps the most crucial prediction of the “molecular-lid” mechanism proposed herein for Conk-C3, is a pivotal role for the toxin Arg54. This prediction is not trivial, as this residue is not conserved among the conkunitzin family (Fig. 1A), and an alanine substitution at the equivalent position of Conk-S1 had no effect on toxicity^20^ (Fig. 6B). The R54A mutant of Conk-C3 blocked Shaker only at extremely high concentrations, corresponding to >300-fold reduction in potency. Differential scattering fluorimetry (DSF) analysis has indicated that this peptide was properly folded (Fig. S5). Despite its marginal activity, Conk-C3 R54A lost its sensitivity to depolarization (Fig. 6C), consistent with the simulated interactions between this toxin residue and the permeant ions at the toxin-channel interface.

## Discussion

Peptide blockers of K^+^ channels bind at their extracellular vestibules and halt ion conduction. Classical blockers that occlude the pore using a conserved lysine residue were extensively employed to study the basic properties of these channels and are being clinically developed for the treatment of various cardiac, neuronal, and autoimmune diseases. Here, we have combined MD simulations with structural and biophysical methods to shed light on a less familiar blocking mechanism that we term a “molecular lid”.

MD simulations suggest three distinct levels of interaction between peptide blockers and K^+^ ions at the selectivity filter (Fig. 5, Movie S3): (*i*) Tight repulsion, in the case of classical blockers such as CTX, prevents ion translocation beyond S2, and increase pore stability; (*ii*) Lose repulsion, in the case of lid-blockers such as Conk-C3, permits ion occupation at all SF sites, but divert ions at the upper SF segment into cryptic coordination sites and induce a gradual widening of the extracellular pore entry; and (*iii*) No interaction – in the case of Conk-S1 an asymmetric collapse of the pore indirectly, but effectively, prevents ion translocation beyond S1.

The “plug” mechanism of classical pore blockers was initially deduced from double-mutant cycle analyses^29^ and trans-enhanced dissociation experiments^27^ and later gained structural support when the CTX-paddle chimera complex structure was solved, revealing an insertion of K27 into the pore and a corresponding missing electron density at the S1 K^+^ coordination site^7^. The “pore-collapse” mechanism proposed for Conk-S1, and observed herein for Conk-C3^RR^ still awaits a direct structural confirmation. Yet, a recent report provides structural evidence that C-type inactivation in K2P2.1 channels entails asymmetric constriction of the SF, which is suppressed by the C-type gate activator ML335^30^. The drug was found to target the peripheral cavities of these channels, in direct support of the MD-based observation that ectopic water traffic at this region tends to destabilize the SF^20,24,31^.

Regarding the “molecular-lid” mode of action proposed here for Conk-C3, there seems to be ample experimental evidence, biophysical and structural, supporting the use of this mechanism by animal toxins. A “lid” or “turret” mode of action was first promoted based on the observation the HERG specific toxin ErgTX blocked currents through this channel despite lacking a conserved lysine, in an [K^+^]_o_ independent manner^18,32^. While the exact molecular details involved in this type of block were not described, several toxins were proposed to share this mode of action based on their biophysical and pharmacological traits^33–37^. The binding of KTX to the KcaA-Kv1.3 chimera was demonstrated to induce the widening of the entrance region to the selectivity filter and rotation of the D80 sidechain, both phenomena interpreted as an induced fit between the toxin and the channel proteins^38^. Here we observed very similar effects of Conk-C3, but not Conk-C3^RR^, during MD simulations of Shaker, which were linked to increased K^+^ occupation at cryptic coordination sites formed between the toxin and the channel molecules.

An alternative binding mode that does not involve a pore-inserting lysine was deduced for Shk-Dap22, an analog with increased selectivity for K_v_1.3, developed as immunosuppressant^39^. Recently, a similar mechanism was identified for the parental toxin and for peptides designed on its scaffold, using phage-display and a novel tethered-toxin methodology^40,41^. Further, the recent suggestion that the classical pore-blocker CTX wobbles between several bound conformations, further imply that the various K^+^ blocking mechanisms portrayed above are not mutually exclusive, and may co-exist, albeit at different potencies, for a given peptide-channel interacting pair.

The last conclusion carries significant implications for the active development of K^+^ blockers as therapeutics, since attempts to improve the pharmacological profile of a given lead molecule by increasing its affinity or by eliminating undesired binding modes may invoke alternative action mechanisms, thus fail to yield the desired results. Observations made in the frame of our previous^20^ and current studies portray MD simulations as a powerful tool to tackle this problem. Simulations of K^+^ channels are highly stable when adequately set and equilibrated, and the molecular events critical for toxin action, i.e., permeation, pore collapse, and cryptic coordination takes place at the sub-microsecond realm. While not identical, trajectories recorded in the presence of a given toxin species successfully encapsulated the key properties associated with its unique mode of action and had a considerable predictive power for the design of subsequent experiments.

**Table 1:** X-ray Data Collection Statistics for the Conk-C1 and Conk-C3 crystals.

|  | **Conk-C1** | **Conk-C3<sup>a</sup>** |
| --- | --- | --- |
| **Data Collection** |  |  |
| PDB code | 6YHT | 6YHY |
| Space group | *P2<sub>1</sub>3* | *P2<sub>1</sub>2<sub>1</sub>2* |
| Cell dimensions: |  |  |
| a,b,c (Å) | 68.61, 68.61, 68.61 | 41.78, 52.28, 56.15 |
| α,β,γ (°) | 90, 90, 90 | 90, 90, 90 |
| No. of copies in a.u. | 1 | 2 |
| Resolution (Å) | 34.30-2.15 | 18.33-1.55 |
| Upper resolution shell (Å) | 2.23-2.15 | 1.56-1.62 |
| Unique reflections | 6072 (603) | 18062 (1700) |
| Completeness (%) | 99.97 (100.00) | 98.35 (94.81) |
| Multiplicity | 10.3 (10.7) | 4.2 (4.3) |
| Average I/σ(I) | 20.58 (1.91) | 17.26 (3.39) |
| R<sub>p</sub>im | 0.01919 (0.3515) | 0.02809 (0.4133) |
| CC1/2 | 0.999 (0.695) | 0.998 (0.587) |
| **Refinement** |  |  |
| Resolution range (Å) | 34.30-2.15 | 18.33-1.55 |
| No. of reflections (I/σ(I) > 0) | 6072 | 18062 |
| No. of reflections in test set | 603 | 1700 |
| R-working / R-free | 0.2050/ 0.2210 | 0.1904/ 0.2245 |
| No. of protein atoms | 437 | 956 |
| No. of water molecules | 10 | 80 |
| Overall average B factor (Å<sup>2</sup>) | 64.58 | 23.31 |
| Root mean square deviations: |  |  |
| - bond length (Å) | 0.007 | 0.005 |
| - bond angle (°) | 0.89 | 0.66 |
| **Ramachandran Plot** |  |  |
| Most favored (%) | 90.74 | 97.37 |
| Additionally allowed (%) | 9.26 | 2.63 |
| Disallowed (%) | 0.0 | 0.0 |

\* Values in parentheses refer to the data of the corresponding upper-resolution shell
<sup>a</sup> The deposited structure contains the 48R/49R substitutions.

## Acknowledgments

We would like to acknowledge prof. Felix Frolow for his invaluable assistance in crystallization and initial data collection of Conk-C1 and Conk-C3.

## Materials and Methods

### Animals

Female wild-caught *Xenopus laevis* frogs were purchased from NASCO. Experiments were performed according to the guidelines of the Weizmann Institute Animal Care and Use Committee permit #37070717-3.

### Method details

#### Expression and purification of Conkunitzin-C1 and Conkunitzin-C3

Gene sequences and native Conkunitzin-C1 and Conkunitzin-C3 were obtained in frame of the CONCO cone snail genome project for health within the 6^th^ Framework Program of the European Commission (a draft genome assembly of the snail is publically available^42^). Synthetic genes encoding the cone peptides were designed applying a codon usage appropriate for bacterial expression. E. coli BL21 (DE3) were transformed with the appropriate toxin construct cloned into the SapI/BamHI sites of pTwin1 (New England Biolabs). Freshly transformed colonies were used to inoculate a 10-ml starter in LB containing 0.1mg/ml Carbenicillin and 0.03mg/ml chloramphenicol. Following 12 hours of growth at 37°C, the starter was diluted into a baffled flask containing 1 liter LB supplemented with antibiotics and grown to OD_600_ of 0.6. IPTG was added to a concentration of 0.4 mM, and culture growth was continued for an additional 5 hours. Cells were harvested by centrifugation (10 min at 5000 × g at 4°C) and re-suspended in 50 ml H2O. The cell suspension was frozen and thawed at 50°C for 45 minutes to disrupt cell walls, and total lysis was achieved by sonication. The cell lysate was centrifuged for 15 min at 20,000 × g and the inclusion bodies pellet was washed and precipitated twice in washing solution containing 25% (W/V) sucrose, 5 mM EDTA, 1% (V/V) Triton X-100, in PBS pH 8.0. The inclusion bodies were dissolved and incubated for 1 hour in 10 ml denaturation buffer containing 6 M Guanidinium-HCl, 0.1 M Tris-HCl pH 8.0, 1 mM EDTA and 10mM reduced glutathione. The denatured protein suspension was diluted 20-fold into cold (4°C) renaturation solution composed of 0.2 M ammonium acetate pH 8.0, 0.5M Sucrose, and 0.2 mM oxidized glutathione and incubated at 4°C with gentle stirring for 24 hours. The renaturation mixture was filtered through 1 mm Whatman paper to eliminate insoluble proteins and the soluble material was precipitated in 50% ammonium sulfate at 4°C for 4 hours. The precipitate was collected by filtration through a GF/C membrane and re-suspended in 20ml phosphate buffer (50mM, pH 6.2) to activate intein cleavage, which releases the toxin from the tag. Following 24 hours incubation at room temperature, the soluble protein was isolated by centrifugation and 0.2μM filtration. The protein sample was then subjected to RP-HPLC, using a 10ml tricorn column packed with source 15RPC on an ACTA-Pure HPLC system. Proteins were eluted with a linear gradient of acetonitrile containing 0.1% TFA (buffer B). Both peptides typically eluted at 23% B, enabling clear separation from the intein moiety that eluted at 70% B. RP-purified samples were dried by lyophilization, re-suspended in water, divided into aliquots, and subsequently dried by speed-vac and stored at −80°C.

#### Structure determination and refinement of the Conk-C1 and Conk-C3 structures

Crystals of Conk-C1 and Conk-C3 (its 48R/49R mutant) were obtained using the hanging-drop vapor-diffusion method. The crystals of Conk-C1 and Conk-C3 were grown from 0.1M Sodium citrate tribasic dihydrate pH 5.6, 20%v/v Isopropanol, and 20% w/v Polyethylene glycol 4,000. The Conk-C1 crystals formed in the cubic space group P213, with one copy per asymmetric unit. A complete dataset, to 2.15 Å resolution was collected at 100 K on a single crystal at the European Synchrotron Radiation Facility (ESRF) beamline, ID29 (wavelength 0.97625 Å). The Conk-C3 crystals formed in the orthorhombic space group P212121, with two copies per asymmetric unit. A complete dataset, to 1.55 Å resolution was collected on beamline ID29. Diffraction images of the Conk-C1 and Conk-C3 crystals were indexed and integrated using the program HKL2000, and integrated intensities were scaled using the program SCALEPACK^43^. The Conk-C1 and Conk-C3 structures were solved by molecular replacement with the program MOLREP^44^.

All steps of atomic refinement of both structures were carried out with the CCP4/REFMAC5 program^45^ and by Phenix refine^46^. The models were built into *2mFobs - DFcalc*, and *mFobs - DFcalc maps* by using the COOT program^47^. Details of the refinement statistics of the Conk-C1 and Conk-C3 structures are described in **Table 1**. The coordinates Conk-C1 and Conk-C3 were deposited in the RCSB Protein Data Bank with accession codes 6YHT and 6YHY, respectively. The structures will be released upon publication.

#### Differential scattering fluorimetry

The thermal stability of Conk-C3 derivatives was determined by differential scanning fluorimetry on a NanoTemper Tycho NT.6 (NanoTemper Technologies, Germany) with a back-reflection aggregation detection at a range from 35 to 95°C and with a heating rate of 20^°^C min^-1^. Protein unfolding at 0.1mM in ND96 buffer was followed by tyrosine fluorescence intensity at 330 and 350 nm. The melting temperature (Tm) was determined by detecting the maximum of the first derivative of the fluorescence ratios (F350/F330) after fitting experimental data with a polynomial function. Data were acquired in triplicates.

#### Electrophysiology – oocytes handling

Plasmids encoding for drosophila ShakerIR and its mutants were linearized using *EcoRI* and purified using phenol/chloroform. The linearized templates were used for mRNA synthesis using the T7 mMESSAGE mMACHINE Transcription Kit (Ambion) and stored as stock solutions at −80°C. *Xenopus laevis* female frogs were anesthetized and subjected to surgery by an incision in the lower half of the belly. The oocytes were pulled out from the incision and placed in a sterile calcium-free ND96 solution containing 96mM NaCl, 2mM KCl, 1mM MgCl_2_ in 5mM HEPES pH 7.5 and then incubated in collagenase solution (3mg/ml) for 2 hours in order to defoliculate them. After the collagenase treatment, the oocytes were incubated in ND96 solution supplemented with 1.8 mM CaCl_2_, 2.5 mM sodium pyruvate, and 100 mg/ml gentamicine (NDE). Selected defolliculated oocytes were injected with 1ng mRNA of interest using a Drummond 510 microdispenser. Injected oocytes were incubated at 18°C for 1-2 days in NDE solution prior to the electrophysiological experiments.

#### Electrophysiology – data acquisition

K^+^ currents from the injected *Xenopus* oocytes were measured by a two-electrode voltage clamp using a Gene Clamp 500 amplifier (Axon Instruments, Union City, CA, USA). Oocytes were placed in a 100 μl fiberglass bath and perfused with ND96. Toxin solutions were freshly made by the re-suspension of lyophilized toxin in ND96 supplemented with 1 mg/ml BSA, and then diluted and applied directly to the bath. Data were sampled at 10 kHz and filtered at 5 kHz using a Digidata 1550A device controlled by pCLAMP 10.5 (Axon Instruments, Union City, CA). Capacitance transients and leak currents were removed by subtracting a scaled control trace utilizing a P/4 protocol.

#### Electrophysiology – Data analysis

**Dose-response curves** were acquired by the application of increasing toxin concentrations into the measurement bath. Each curve was constructed from at least 5 toxin concentrations, an adequate period of incubation (> 1 min) at each toxin concentration ensured a steady-state response. Experiments were carried in triplicates. Data points were fit using a Hill equation with the hill coefficient restricted to unity:

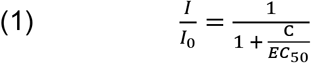

Where I_0_ is the unmodified current measured before toxin application, C is toxin concentration, and EC_50_ is the half-maximal toxin concentration.

**Voltage-dependent toxin dissociation assays** - Oocytes expressing Shaker were incubated with a pre-set toxin dose for 5 minutes until a stable block of ~80% was achieved. Voltage-dependent toxin-dissociation was assayed using a two-pulse voltage protocol (Fig. 2C, inset) in which two test pulses (P1,P2) of 50ms to −10mV applied from a holding potential of −90mV were separated by a strong depolarizing pulse (+100mV) of variable duration. Voltage-dependent dissociation of the bound toxin leads to an elevated amplitude of the current recorded during the second test pulse and an increase of the ratio P2/P1.

### MD simulations

#### Modeling of the Shaker-Conk-C3 and Shaker-Conk-C3^RR^ complexes

A structural model of the Shaker pore domain (residues Lys376-Asp488) was constructed based on the published crystal structure of the Kv1.2-Kv2.1 paddle chimera (PDB ID 2R9R, 86% identity, 94% similarity) using MODELER^48^. For docking Conk-C3^RR^, the 1.55Å crystal structure of this mutant (PDB ID: 6YHY) was used; the structure of the wild-type toxin was produced by applying the Rotamer tool in UCSF chimera to substitute R48 and R49 of Conk-C3^RR^ back to Ile and His that occupy these positions in the original sequence. The resulting models were submitted for a rigid-body docking without any constraints using the CLUSPRO web service ^49^, and the solutions were ranked using the weighting coefficients of the Van der Waals+electrostatics docking mode. For both toxin derivatives, the majority of the top 1000 solutions were divided between four clusters, which could be converged into a single unique solution by rotation about the pore axis.

#### MD Simulation protocol

The simulation systems employed in this study were assembled using the CHARMM-GUI web service^50^. The channel protein was embedded in a lipid bilayer roughly 100Å^2^ in size composed of a 3:1 POPC:POPG mixture, and the water thickness was set to 22.5 Å. Two K^+^ ions were initially placed at sites s2, and s4 and a water molecule was placed at site s3 of the selectivity filter based on the crystallographic coordinates to obtain an initial 2,4 configuration. Additional K^+^ and Cl^-^ ions were added to neutralize the system and set the KCl concentration at 150mM. We have used the CHARMM36m parameter set for the various system components^50^ and the TIP3P model for water. Simulations were carried on GPU using GROMACS 2019^51^. All simulations were performed under NPT conditions at 303K and 1atm. Periodic boundary conditions and electrostatic interactions were treated using the particle mesh Ewald method. A time step of 2fs was employed, and the hydrogen atoms were constrained using LINCS. Assembled systems were minimized and equilibrated using gradual melting of sidechain and backbone atoms^52^, and an unbiased MD run of 20ns was carried as the final equilibration step.

#### Analysis protocols for MD simulations

Trajectory snapshots were saved every 100ps during production simulations. A custom written MATLAB code was used for trajectory analysis. The peripheral cavity of a given channel subunit was empirically defined as the volume formed by the intersection of four spheres, with centers coordinates at the backbone nitrogen atoms of Val438, Tyr445, Met448, and Val451 and with radii of 10, 7, 10 and 14Å respectively. This definition effectively filtered out pore-waters and water molecules from the neighboring cavities.

#### Quantification and Statistical Analysis

All statistical analyses were conducted using MATLAB built-in functions. All data are expressed as the mean ± standard deviation unless otherwise stated in the figure legend.

## Supplementary Figure Legends

**Figure S1.**
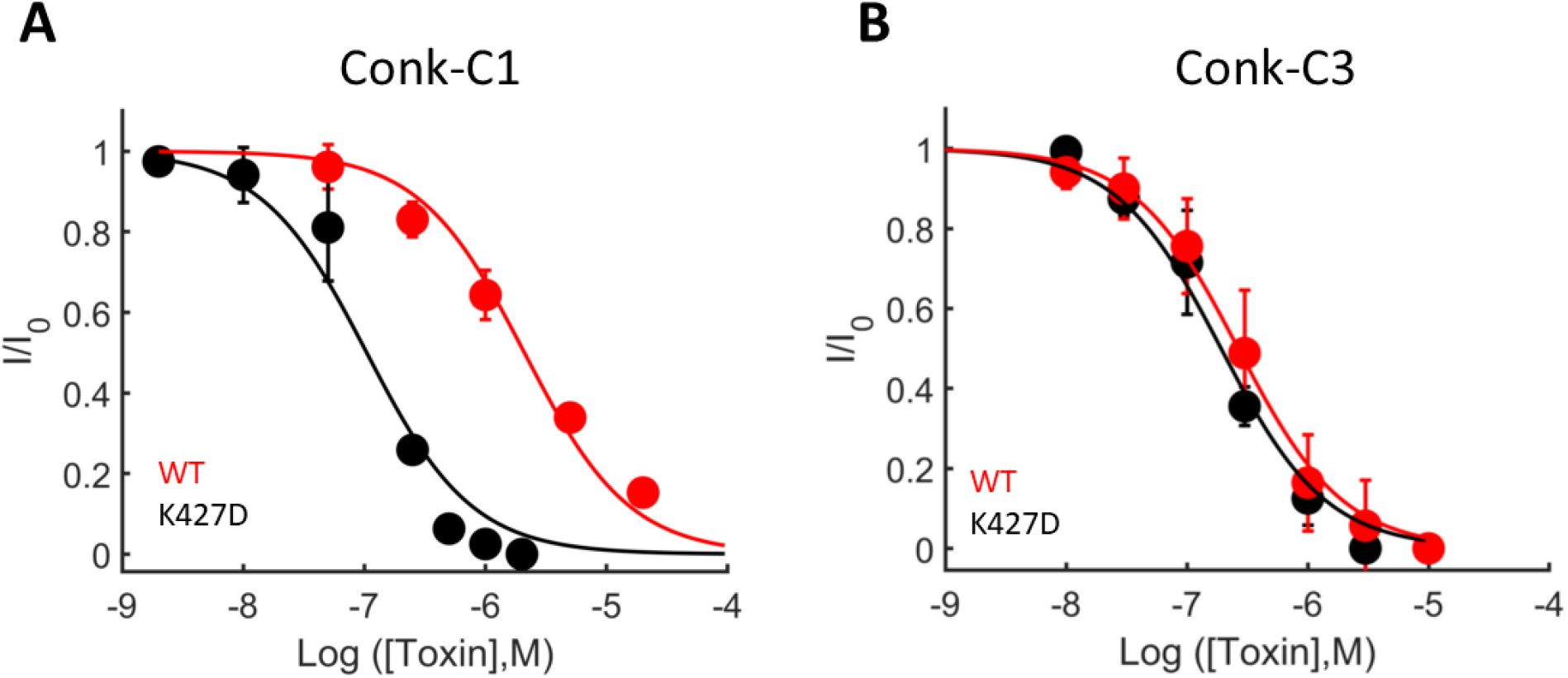
Dose-response curves of Conk-C1 (left) and Conk-C3 (right) on wild type Shaker (black) and the K427D mutant (red). Determinations were carried in triplicates; error bars represent standard deviations. EC_50_ values determined were 2.2±0.1μM and 166±31nM on the wild-type shaker, and 108±18nM, 190±27nM on Shaker_KD_ for Conk-C1 and Conk-C3 respectively.

**Figure S2.**
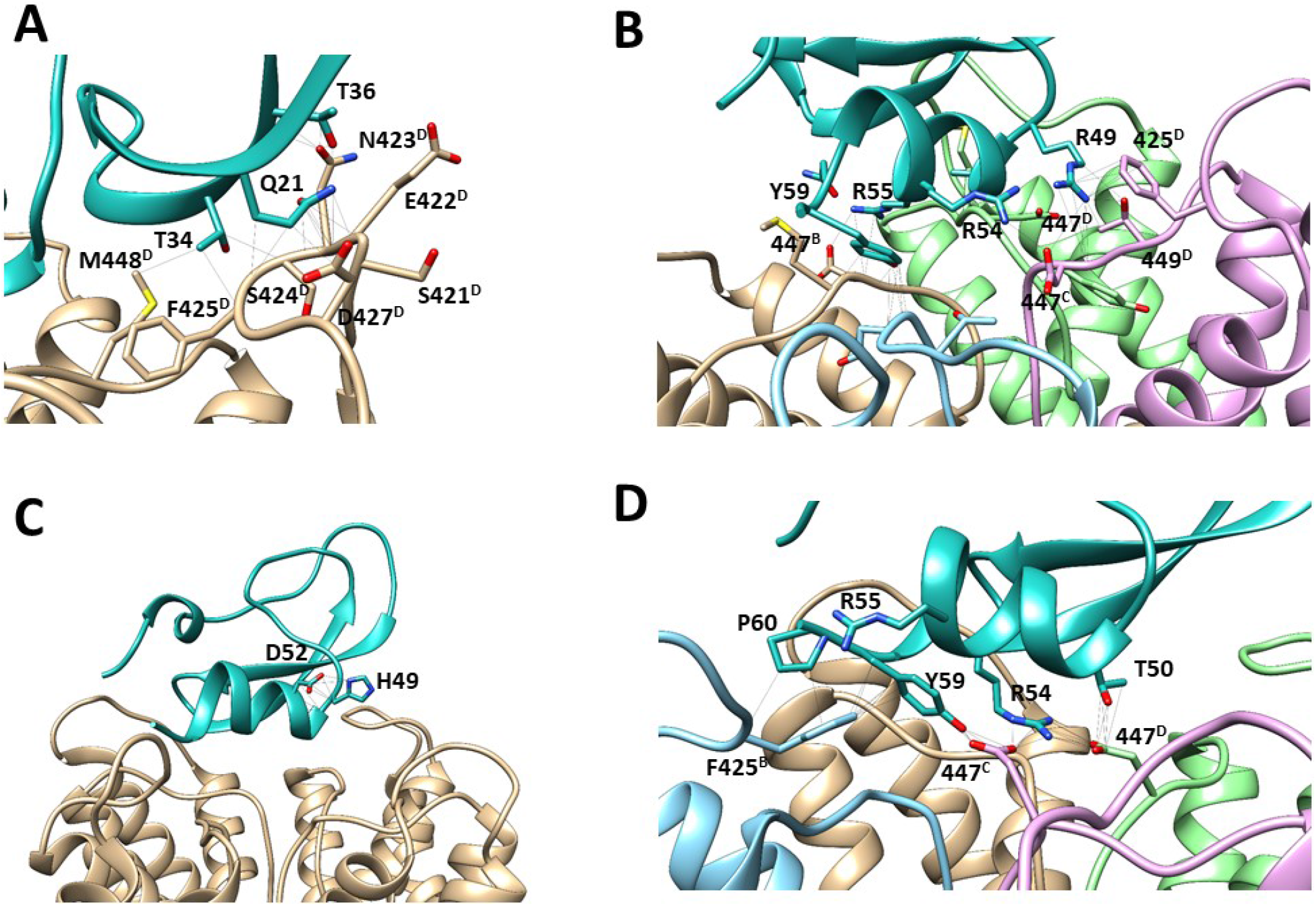
Comparison of protein-protein interactions in the bound poses of Conk-C3^RR^ (top) and the wild-type Conk-C3 (bottom). Channel residues are denoted using a residue^chain^ notation.

**Figure S3.**
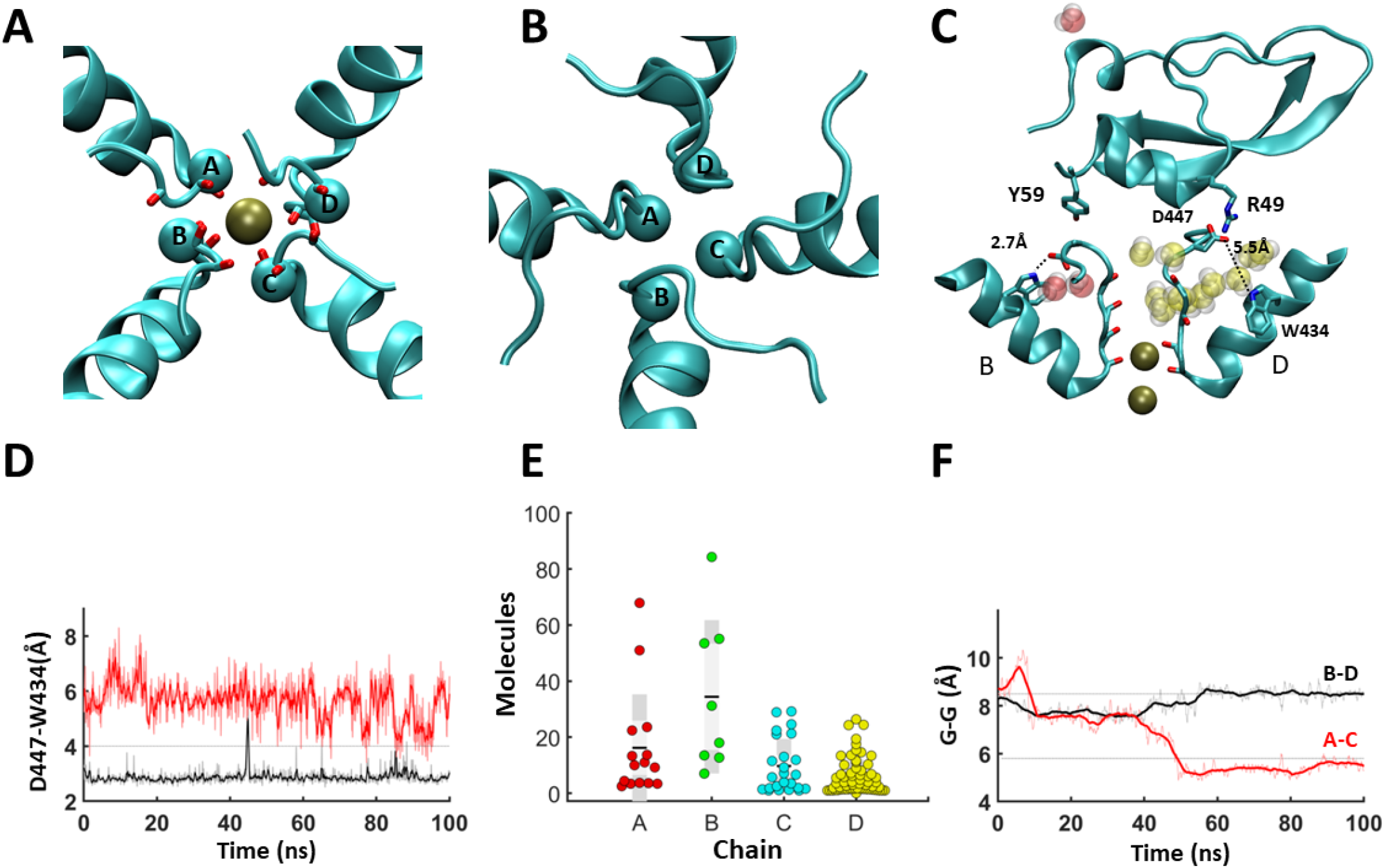
A, Asymmetric dilation observed at the upper segment of the pore in simulations of bound Conk-C3. The snapshot was taken at t=100ns, the Cα atoms of Tyr445 are depicted as spheres, the diagonals were AC = 8.4Å, BD = 10.0Å. B, Asymmetric constriction of the SF during MD simulation of bound Conk-C3^RR^. The snapshot was taken at t=100ns, spheres represent the Cα atoms of Gly444, with measured diagonals of AC=5.5Å, BD=8.5Å. C, side-view of the pore region, and the bound toxin from the snapshot depicted in B. Water molecules assigned to the peripheral pockets of chain B (red) and D (yellow) during the run are depicted as transparent spheres. Key interactions made by the toxin molecule and the apertures of the corresponding DW gates are indicated. D-F, DW apertures (D), Water traffic (E), and pore symmetry (F) dynamics during the trajectory depicted in panels B,C.

**Figure S4.**
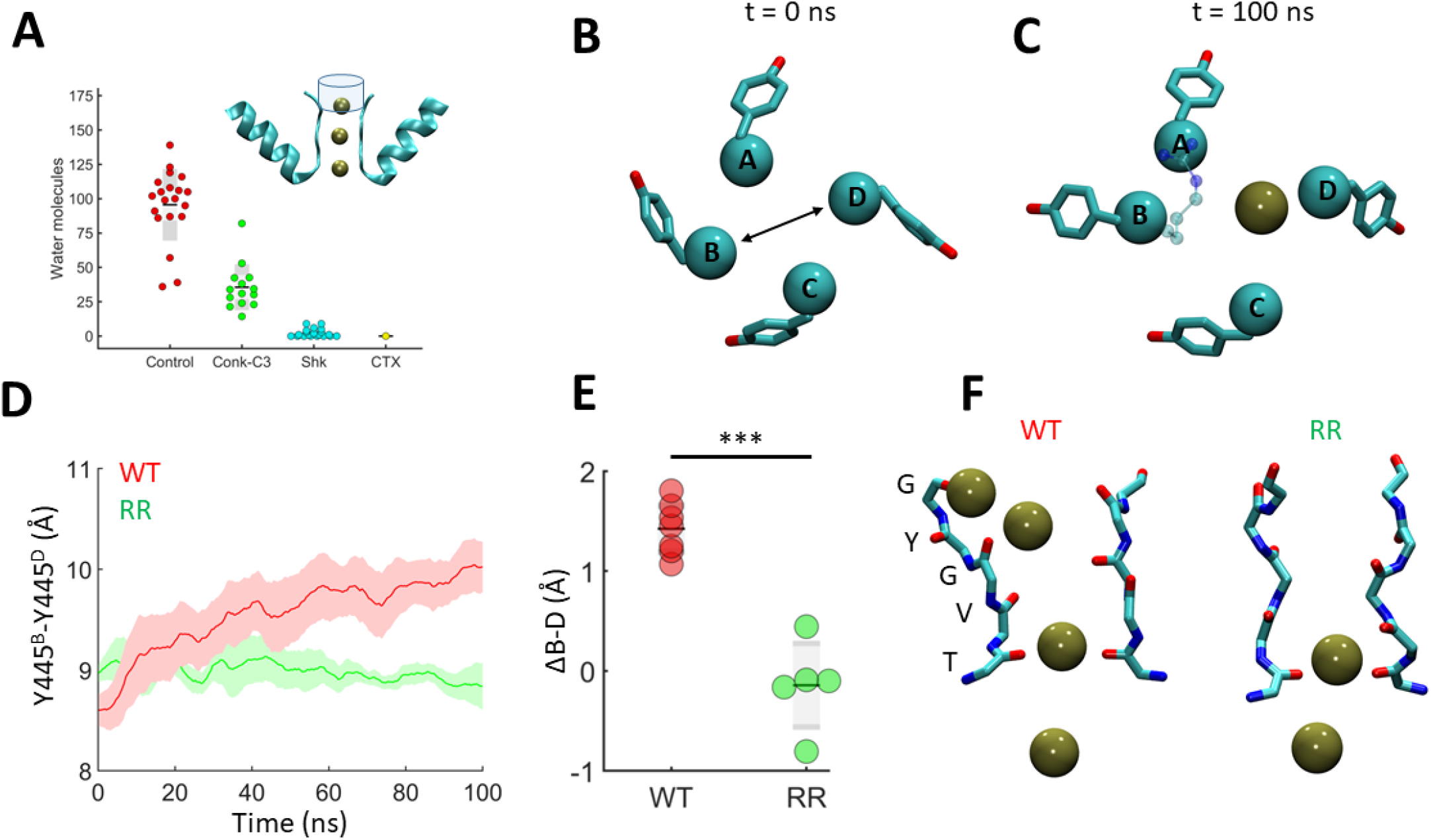
A, The number of water molecules passing through a cylindrical volume positioned just above the SF (inset) during a 10ns simulation window of unmodified (red), Conk-C3 (green), Shk (cyan) and CTX (yellow) - bound systems. Control Data for CTX and Shk were presented in a previous report^20^ and represent a single 200ns trajectory. For Conk-C3, each point represents a discrete 100ns trajectory (14 trajectories in total). B, C, Dilation at the top SF segment in the presence of Conk-C3 observed following electric field (EF) application. The ring of Tyr445 residues is presented with Cα atoms rendered in space fill. B, The Cα - Cα distance between chains B,D captured in panels D, and E is indicated. C, K^+^ Ion bound in S0 is rendered as a solid sphere, Conk-C3 R54 sidechain is overlayed in transparent balls and sticks. D, Pore-widening in the B-D axis during EF simulations of Conk-C3 (red) and Conk-C3^RR^ (green). The shown traces were averaged over 7 and 5 simulations for Conk-C3 and Conk-C3^RR^, respectively, shaded areas represent STD. E, Comparison of the widening at the B-D axis taken as the difference between this distance at the beginning and the end of each simulation reported in (D). F, typical ionic configurations at the SF during EF simulations of Conk-C3 (red) and Conk-C3^RR^ (green) bound systems. SF residues are designated, a kink resulting from an asymmetric collapse of the SF appears at S2 in the Conk-C3^RR^ – bound systems.

**Figure S5.**
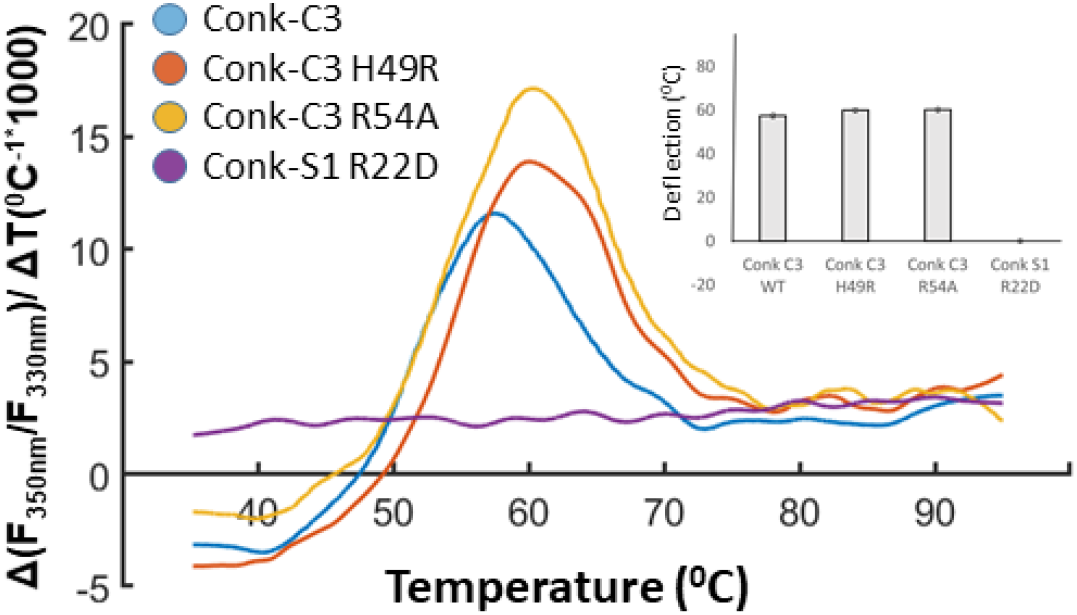
First derivative of the ratio F_350nm_/F_330nm_ representing shifts of the tyrosine fluorescence as a function of temperature. Curve peaks represent the deflection points of the various toxin mutants. (Colors: blue – Conk-C3, orange – H49R, yellow-R54, and purple – Conk-S1 R22D). Conk-S1 R22D is known to misfold due to the elimination of essential intramolecular packing interactions. Inset: deflection temperature calculated for the various mutants, n=3, error bars are S.E.M.

**Movie S1.**

Repulsion between Conk-C3:R54 and a K^+^ ion during an outward permeation transition. The movie depicts a 30ns window in which a knock-in event takes place, and a K^+^ ion is promoted from S2 to S0. Conk-C3:R54 is depicted in yellow, and its displacements from Y445 and G446 carbonyls as well as D447^A^:Oδ_2_ (Fig. 4A a,b,c) are indicated.

**Movie S2.**

A cryptic K^+^ coordination site formed at the interface between chains C and D of the channel and the toxin molecule. The movie span a 50ns time window in which an S2->S0 transition takes place. The events described in Fig. 3 B, C are taking place at t=10 and t=32 ns, respectively.

**Movie S3.**

Three distinct levels of interaction between peptide blockers and K^+^ ions at the selectivity filter. K^+^ dynamics at the SF along a 50ns trajectory is compared between free Shaker_KD_ and channel bound to CTX, Conk-S1 or Conk-C3. Two opposing channel subunits are shown, for each toxin residues critical for action are depicted.

## References

1. Hille, B. Ion Channels of Excitable Membranes, 3rd edition. (2001).

2. Kuang, Q., Purhonen, P. & Hebert, H. Structure of potassium channels. Cell. Mol. Life Sci. 72, 3677–3693 (2015).

3. Heginbotham, L., Lu, Z., Abramson, T. & MacKinnon, R. Mutations in the K+ channel signature sequence. Biophys. J. 66, 1061–1067 (1994).

4. Doyle, D. a. et al. The Structure of the Potassium Channel: Molecular Basis of K+ Conduction and Selectivity. Science 280, 69–77 (1998).

5. Zhou, Y., Morais-Cabral, J. H., Kaufman, A. & Mackinnon, R. Chemistry of ion coordination and hydration revealed by a K+channel-Fab complex at 2.0 Å resolution. Nature 414, 43–48 (2001).

6. Kalia, J. et al. From foe to friend: Using animal toxins to investigate ion channel function. J. Mol. Biol. 427, 158–175 (2015).

7. Banerjee, a., Lee, a., Campbell, E. & MacKinnon, R. Structure of a pore-blocking toxin in complex with a eukaryotic voltage-dependent K+ channel. Elife 2, e00594–e00594 (2013).

8. Park, C. S. & Miller, C. Interaction of charybdotoxin with permeant ions inside the pore of a K+ channel. Neuron 9, 307–13 (1992).

9. Lange, A. et al. Toxin-induced conformational changes in a potassium channel revealed by solid-state NMR. Nature 440, 959–62 (2006).

10. Rauer, H., Pennington, M., Cahalan, M. & Chandy, K. G. Structural conservation of the pores of calcium-activated and voltage-gated potassium channels determined by a sea anemone toxin. J. Biol. Chem. 274, 21885–21892 (1999).

11. Chandy, K. G. et al. K+ channels as targets for specific immunomodulation. Trends Pharmacol. Sci. 25, 280–289 (2004).

12. Murray, J. K. et al. Pharmaceutical Optimization of Peptide Toxins for Ion Channel Targets: Potent, Selective, and Long-Lived Antagonists of Kv1.3. J. Med. Chem. 58, 6784–6802 (2015).

13. Kalman, K. ShK-Dap22, a Potent Kv1.3-specific Immunosuppressive Polypeptide. J. Biol. Chem. 273, 32697–32707 (1998).

14. Mouhat, S. et al. Pharmacological profiling of Orthochirus scrobiculosus toxin 1 analogs with a trimmed N-terminal domain. Mol. Pharmacol. 69, 354–362 (2006).

15. Pennington, M. W. et al. Engineering a Stable and Selective Peptide Blocker of the Kv1.3 Channel in T Lymphocytes. Mol. Pharmacol. 75, 762–773 (2009).

16. Beeton, C. et al. Targeting effector memory T cells with a selective peptide inhibitor of Kv1.3 channels for therapy of autoimmune diseases. Mol. Pharmacol. 67, 1369–1381 (2005).

17. Gordon, D., Chen, R. & Chung, S. H. Computational methods of studying the binding of toxins from venomous animals to biological ion channels: Theory and applications. Physiol. Rev. 93, 767–802 (2013).

18. Giaever, G., Xu, C.-Q., Zhu, S.-Y., Chi, C.-W. & Tytgat, J. Turret and pore block of K+ channels: what is the difference? Trends Pharmacol. Sci. 24, 446–448 (2003).

19. Rodríguez De La Vega, R. C., Merino, E., Becerril, B. & Possani, L. D. Novel interactions between K+channels and scorpion toxins. Trends in Pharmacological Sciences vol. 24 222–227 (2003).

20. Karbat, I. et al. Pore-modulating toxins exploit inherent slow inactivation to block K+ channels. Proc. Natl. Acad. Sci. 116, 18700–18709 (2019).

21. Violette, A. et al. Large-scale discovery of conopeptides and conoproteins in the injectable venom of a fish-hunting cone snail using a combined proteomic and transcriptomic approach. J. Proteomics 75, 5215–25 (2012).

22. Bayrhuber, M. et al. Conkunitzin-S1 is the first member of a new Kunitz-type neurotoxin family. Structural and functional characterization. J. Biol. Chem. 280, 23766–70 (2005).

23. Baştuğ, T. & Kuyucak, S. Importance of the peptide backbone description in modeling the selectivity filter in potassium channels. Biophys. J. 96, 4006–12 (2009).

24. Li, J., Ostmeyer, J., Cuello, L. G., Perozo, E. & Roux, B. Rapid constriction of the selectivity filter underlies C-type inactivation in the KcsA potassium channel. J. Gen. Physiol. 150, (2018).

25. Childers, M. C. & Daggett, V. Validating Molecular Dynamics Simulations against Experimental Observables in Light of Underlying Conformational Ensembles. J. Phys. Chem. B 122, 6673–6689 (2018).

26. Hollingsworth, S. A. & Dror, R. O. Molecular Dynamics Simulation for All. Neuron 99, 1129–1143 (2018).

27. Goldstein, S. a & Miller, C. Mechanism of charybdotoxin block of a voltage-gated K+ channel. Biophys. J. 65, 1613–9 (1993).

28. Moldenhauer, H., Díaz-Franulic, I., Poblete, H. & Naranjo, D. Trans-toxin ion-sensitivity of charybdotoxin-blocked potassium-channels reveals unbinding transitional states. Elife 8, 1–23 (2019).

29. Hidalgo, P. & MacKinnon, R. Revealing the architecture of a K+ channel pore through mutant cycles with a peptide inhibitor. Science (80-.). 268, 307–310 (1995).

30. Lolicato, M. et al. K_2P_ channel C-type gating involves asymmetric selectivity filter order-disorder transitions. bioRxiv 2020.03.20.000893 (2020) doi:10.1101/2020.03.20.000893.

31. Ostmeyer, J., Chakrapani, S., Pan, A. C., Perozo, E. & Roux, B. Recovery from slow inactivation in K+ channels is controlled by water molecules. Nature 501, 121–4 (2013).

32. Pardo-Lopez, L. et al. Mapping the binding site of a human ether-a-go-go-related gene-specific peptide toxin (ErgTx) to the channel’s outer vestibule. J. Biol. Chem. 277, 16403–16411 (2002).

33. Zhang, M. et al. BeKm-1 is a HERG-specific toxin that shares the structure with ChTx but the mechanism of action with ErgTx1. Biophys. J. 84, 3022–3036 (2003).

34. Al-Sabi, A. et al. κM-conotoxin RIIIK, structural and functional novelty in a K + channel antagonist. Biochemistry 43, 8625–8635 (2004).

35. Finol-Urdaneta, R. K. et al. Marine toxins targeting KV1 channels: Pharmacological tools and therapeutic scaffolds. Mar. Drugs 18, (2020).

36. M’Barek, S. et al. Synthesis and characterization of Pi4, a scorpion toxin from Pandinus imperator that acts on K+ channels. Eur. J. Biochem. 270, 3583–3592 (2003).

37. Mouhat, S. et al. The functional dyad of scorpion toxin Pi1 is not itself a prerequisite for toxin binding to the voltage-gated Kv1.2 potassium channels. Biochem. J. 377, 25–36 (2004).

38. Zachariae, U. et al. The molecular mechanism of toxin-induced conformational changes in a potassium channel: relation to C-type inactivation. Structure 16, 747–54 (2008).

39. Lanigan, M. D. et al. Mutating a critical lysine in ShK toxin alters its binding configuration in the pore-vestibule region of the voltage-gated potassium channel, Kv1.3. Biochemistry 41, 11963–71 (2002).

40. Zhao, R., Dai, H., Mendelman, N., Chill, J. H. & Goldstein, S. A. N. Tethered peptide neurotoxins display two blocking mechanisms in the K+ channel pore as do their untethered analogs. Sci. Adv. 6, eaaz3439 (2020).

41. Zhao, R. et al. Designer and natural peptide toxin blockers of the KcsA potassium channel identified by phage display. Proc. Natl. Acad. Sci. U. S. A. 112, E7013–E7021 (2015).

42. Andreson, R. et al. Gene content of the fish-hunting cone snail Conus consors. bioRxiv 590695 (2019) doi:10.1101/590695.

43. Otwinowski, Z. & Minor, W. B. T.-M. in E. [20] Processing of X-ray diffraction data collected in oscillation mode. in Macromolecular Crystallography Part A vol. 276 307–326 (Academic Press, 1997).

44. Vagin, A. & Teplyakov, A. Molecular replacement with MOLREP. Acta Crystallogr. D. Biol. Crystallogr. 66, 22–25 (2010).

45. Murshudov, G. N., Vagin, A. A. & Dodson, E. J. Refinement of Macromolecular Structures by the Maximum-Likelihood Method. Acta Crystallogr. Sect. D 53, 240–255 (1997).

46. Afonine, P. V et al. Towards automated crystallographic structure refinement with phenix.refine. Acta Crystallogr. Sect. D 68, 352–367 (2012).

47. Emsley, P. & Cowtan, K. Coot: Model-building tools for molecular graphics. Acta Crystallogr. Sect. D Biol. Crystallogr. 60, 2126–2132 (2004).

48. Webb, B. & Sali, A. Protein Structure Modeling with MODELLER BT - Functional Genomics: Methods and Protocols. in (eds. Kaufmann, M., Klinger, C. & Savelsbergh, A.) 39–54 (Springer New York, 2017). doi:10.1007/978-1-4939-7231-9_4.

49. Kozakov, D. et al. The ClusPro web server for protein-protein docking. Nat. Protoc. 12, 255–278 (2017).

50. Jo, S., Kim, T., Iyer, V. G. & Im, W. CHARMM-GUI: A web-based graphical user interface for CHARMM. J. Comput. Chem. 29, 1859–1865 (2008).

51. Abraham, M. J. et al. GROMACS: High performance molecular simulations through multi-level parallelism from laptops to supercomputers. SoftwareX 1–2, 19–25 (2015).

52. Wu, E. L. et al. CHARMM-GUI Membrane Builder toward realistic biological membrane simulations. J. Comput. Chem. 35, 1997–2004 (2014).

53. Hu, H., Bandyopadhyay, P. K., Olivera, B. M. & Yandell, M. Characterization of the Conus bullatus genome and its venom-duct transcriptome. BMC Genomics 12, (2011).

54. Pardos-Blas, J. R., Irisarri, I., Abalde, S., Tenorio, M. J. & Zardoya, R. Conotoxin diversity in the venom gland transcriptome of the magician’s cone, pionoconus magus. Mar. Drugs 17, (2019).

55. Gasparini, S. et al. Delineation of the functional site of α-dendrotoxin: The functional topographies of dendrotoxins are different but share a conserved core with those of other Kv1 potassium channel-blocking toxins. J. Biol. Chem. 273, 25393–25403 (1998).

56. Huber, R. et al. Structure of the complex formed by bovine trypsin and bovine pancreatic trypsin inhibitor. II. Crystallographic refinement at 1.9 Å resolution. J. Mol. Biol. 89, 73–101 (1974).

